# The suppressive cap-binding-complex factor 4EIP is required for normal differentiation

**DOI:** 10.1101/314997

**Authors:** Monica Terrao, Kevin Kamanyi Marucha, Elisha Mugo, Dorothea Droll, Ihor Minia, Franziska Egler, Johanna Braun, Christine Clayton

## Abstract

*Trypanosoma brucei* live in mammals as bloodstream forms and in the Tsetse midgut as procyclic forms. Differentiation from one form to the other proceeds via a growth-arrested stumpy form with low mRNA content and translation. The parasites have six eIF4Es and five eIF4Gs. EIF4E1 pairs with the mRNA-binding protein 4EIP but not with any EIF4G. EIF4E1 and 4EIP each inhibit expression when tethered to a reporter mRNA. The 4E-binding motif in 4EIP is required for the interaction with EIF4E1 both *in vivo* and in a 2-hybrid assay, but not for the suppressive activity of 4EIP when tethered. However, the suppressive activity of EIF4E1 when tethered requires 4EIP. Correspondingly, in growing bloodstream forms, 4EIP is preferentially associated with unstable mRNAs. Trypanosomes lacking 4EIP have a marginal growth disadvantage as cultured bloodstream or procyclic forms. Bloodstream forms without 4EIP cannot make differentiation-competent stumpy forms, but the defect can be complemented by a truncated 4EIP that does not interact with EIF4E1. Bloodstream forms lacking EIF4E1 have a growth defect but can differentiate. We suggest that 4EIP and EIF4E1 fine-tune mRNA levels in growing cells, and that 4EIP is required for mRNA suppression during differentiation to the stumpy form.

## Introduction

The amounts of protein synthesised from an mRNA are determined by the rates of translation initiation (1,2) and elongation (3,4). Most eukaryotic translation is initiated by binding of eIF4E to the cap structure. eIF4E recruits eIF4G, which in turn recruits the helicase eIF4A and, via other initiation factors, the 43S complex. The 43S complex, which includes the 40S subunit with charged methionyl tRNA as well as various additional translation factors, then scans towards the start codon (5). Most eukaryotic species examined have more than one eIF4E homologue (6). The roles of different eIF4Es have been most extensively investigated in metazoa; in general there are one or two eIF4Es that are responsible for constitutive cap-dependent translation and the remainder have specialised roles at particular developmental stages, or are involved in processes other than translation (6).

In animal cells, one mechanism for the regulation of translation initiation involves 4E binding proteins (4EBPs) (7). The 4E-BP - eIF4E interaction is partially mediated by a canonical binding motif, YXXXXLØ, which is found near 4E-BP N-termini (8). Since eIF4G binds eIF4E using the same motif, 4EBP and eIF4G binding is mutually exclusive. 4E-BP - eIF4E binding is however strengthened by additional interactions (9-11).

The “tethering” assay is a method that measures the effects of attachment of a protein to a reporter mRNA. The protein of interest is expressed as a fusion with a protein or peptide that has the ability to bind a short RNA sequence with high affinity and specificity, such as the lambdaN peptide. Activity is monitored using a reporter mRNA that includes the cognate RNA sequence - for the lambdaN peptide, boxB. Tethering of various 4E-BPs results in translation repression but curiously, this does not require the YXXXXLØ motif. For example, the abilities of tethered 4E-T or *Drosophila* Cup to suppress a target reporter mRNA does not depend on YXXXXLØ (12,13). In both cases, the authors suggested that the tethered protein was able to recruit the CAF1-NOT deadenylase complex (12,13). In contrast to the tethering results, exogenous expression of mammalian 4E-T resulted in indiscriminate translational suppression, and this was dependent on the YXXXXLØ motif (13), suggesting recruitment of 4E-T via eIF4E. It is not known whether tethered 4E-BPs without YXXXXLØ can bind to eIF4E *in vivo* using their additional binding surface (9-11), or whether they act independently of eIF4E.

Another mechanism by which translation can be inhibited in animals and plants depends on eIF4E-like cap-binding proteins that cannot recruit eIF4G. These include mammalian 4EHP/eIF4E2 (14), *Drosophila* d4EHP/EIF4E-8 (15,16), and Arabidopsis nCBP (17). Mammalian and *Drosophila* 4EHPs are required for correct translation control during development (18-21). Interactions with partner proteins such as Bicoid (16) or GIGYF2 (=GIF2) (22-24), and with the RISC complex (25,26), have been implicated in induction of mRNA decay and/or translation suppression by 4EHPs. GIGYF2 appears to be able to act both in conjunction with (24), and independently of, 4EHP (27), and interactions with both Ago2 (28) and the CAF1/NOT complex (27) have been documented.

*Trypanosoma brucei* is a unicellular eukaryotic parasite. The bloodstream form lives extracellularly in the blood and tissue fluids of mammals, evading the immune response by antigenic variation of a variant surface glycoprotein coat (29). The parasites multiply as long slender bloodstream forms, which depend on glycolysis for their energy metabolism (30). When parasitaemia reaches a critical level, non-dividing “stumpy forms” start to dominate the population (31,32). Although these cells have very low overall mRNA and translation levels (33,34), they do express a variety of new proteins, including specific markers such as the surface protein PAD1 (35), and some proteins of mitochondrial energy metabolism (31). Stumpy forms are pre-adapted for development in the definitive host, the Tsetse fly, which becomes infected by blood feeding. Within the Tsetse midgut the parasites differentiate to procyclic forms, which have EP and GPEET procyclins on their surface (36) and a mitochondrial energy metabolism that depends on amino acid substrates (37-39). After a few weeks, with additional differentiation steps, mammal-infective parasites reappear in the salivary glands and can be transmitted at the next blood meal (40).

Cellular differentiation requires substantial changes in gene expression, which usually depend on regulatory transcription factors. Trypanosomes and related organisms, however, lack transcriptional control of individual genes. Instead, transcription is polycistronic: individual mRNAs are excised by processing (41). The 5’-end of each mRNA is formed by *trans* splicing of a 39nt leader sequence (42), which bears a highly methylated cap called cap4: m^7^Gpppm_3_^6,6,2’^ Apm^2^ Apm^2^ Cpm_2_^3,2^ U (43,44). The 3’-end is created by cleavage and polyadenylation (45,46). As a consequence of this arrangement, regulation of gene expression in trypanosomes is almost exclusively post-transcriptional: there are big differences between mRNAs in their processing efficiencies, decay rates, and translation (47-49). RNA-binding proteins have been shown to regulate specific trypanosome mRNA subsets, acting on splicing, mRNA decay, and / or translation (50-52). With one exception, however (53), the molecular mechanisms by which these proteins act are not yet known.

This paper concerns control of translation in *T. brucei*. *T. brucei* and the closely related parasite *Leishmania* have six eIF4Es and five eIF4Gs (54). Results from studies of protein-protein interactions, migration in sucrose gradients, intracellular location, and the effects of RNAi in *T. brucei* suggest that the complexes of eIF4E4 with eIF4G3, and of EIF4E3 with EIF4G4, might be implicated in general translation initiation (54). EIF4E3 and EIF4E4 are present in roughly equimolar ratios, or a slight excess, relative to mRNA (55,56). RNAi experiments showed that EIF4E3 is essential in both forms whereas for EIF4E4, a growth defect was seen only in bloodstream forms (57). The *T. brucei* EIF4E5-EIF4G2, EIF4E5-EIF4G1 (58), and EIF4E6-EIF4G5 (59) complexes might be implicated in translation of specific mRNA subsets, but these have yet to be characterised; alternative functions are also conceivable. *T. brucei* eIF4E2 is not essential in procyclic forms; it interacts with an RNA-binding protein, SLBP2, which is similar to mammalian histone-mRNA-binding proteins (60).

EIF4E1 has been studied most extensively in *Leishmania major*. It binds cap4 and m^7^GTP with a relatively low affinity which is, however, only 3 times worse than that of *Leishmania* EIF4E4 (61,62). *L. major* EIF4E1 does not interact with any of the five eIF4G homologues, either by pull-down or by yeast 2-hybrid analysis (63). In contrast, it does interact with another protein, 4EIP (63), and also directly with EIF3a (64). The latter interaction could result in EIF4G-independent translation initiation. Because of life-cycle-dependent differences in both the EIF4E1-4EIP interaction, and in expression of other EIF4Es, it was suggested the EIF4E1 might act as an unconventional initiation factor in the intracellular amastigote life-cycle stage. Further, it was hypothesised that the initiation activity of EIF4E1 is suppressed, in the promastigote stage, by interaction with 4EIP (63). The results of yeast two-hybrid assays indicated that a YXXXXLØ motif at the N-terminus of *Leishmania* 4EIP was required for interaction with EIF4E1 (63), and co-crystallization revealed that first 52 residues of 4EIP form two alpha-helices that interact with EIF4E1; the first includes the consensus motif (65). *In vitr*o evidence suggested that binding of m^7^GTP by EIF4E1 was inhibited by addition of the N-terminal 4EIP fragment, probably as a consequence of a conformational change within the cap-binding pocket (65).

*T. brucei* EIF4E1 (Tb927.11.2260) is found in the nucleus and cytosol (55). Since there are 3 times fewer EIF4E1 molecules per cell than there are mRNAs (55,66), it cannot be a major general translation initiation factor. Depletion of *T. brucei* eIF4E1 by RNAi halted growth of bloodstream forms and slowed growth of procyclic forms (57). EIF4E1 and 4EIP (Tb927.9.11050) were found to be extremely strong repressors when tethered to reporter mRNAs (67,68). We therefore decided to investigate the function of 4EIP in *T. brucei*. We were particularly interested in knowing whether 4EIP is recruited to specific mRNAs via RNA-binding proteins, and also in knowing the role of 4EIP during differentiation.

## Results

### 4EIP sequence comparisons

To obtain preliminary indications concerning 4EIP functional conservation, we searched for similar sequences and found potential homologues in many kinetoplastid genomes, including that of the free-living *Bodo caudatus*. (The absence of 4EIP in some kinetoplastid genomes may be explained by the fact that not all sequences are complete.) The N-terminal 4E interaction domain, YxxxxLØ, is found in all complete sequences (Fig 1A and Supplementary Fig S1). All but one of the hydrophobic residues in the second EIF4E1-interacting helix (65) are also conserved, suggesting that they are important for 4EIP function. A second, highly charged, conserved region (Fig 1A and Supplementary Fig S1) is separated from the EIF4E1 interaction domain by disordered linker. In *B. caudatus*, *L. major*, *Trypanosoma theileri* and *Trypanosoma grayi*, the linker includes polyglycine tracts (Supplementary Fig S1). The remainder of the protein is much less conserved, except that the C-termini are proline- and glutamine-rich, and highly basic. The last 183 residues of *T. brucei* 4EIP have 21% glutamine and 17% proline and a predicted pI of 10.8. Values for the equivalent regions from *Bodo caudatus* and *Endotrypanum* are, respectively, 39% and 21% glutamine; 11% and 20% proline; and pIs of 11.5 and 12.0. All of the 4EIPs listed in Fig 1 have polyglutamine repeats at the far C-terminus, with the exception of those from the salivarian trypanosomes (*congolense*, *brucei* and *vivax*). The complete sequences terminate with (Q)_1-17_RR, except in *Bodo* which has (Q)_11_R. There is no indication from the sequences of which part might be implicated in mRNA binding, although the C-termini may bind via charge alone. *T. brucei* 4EIP is predicted by Phyre (69) to be predominantly disordered (Supplementary Fig S1, pages 2 & 3).

**Figure 1.**
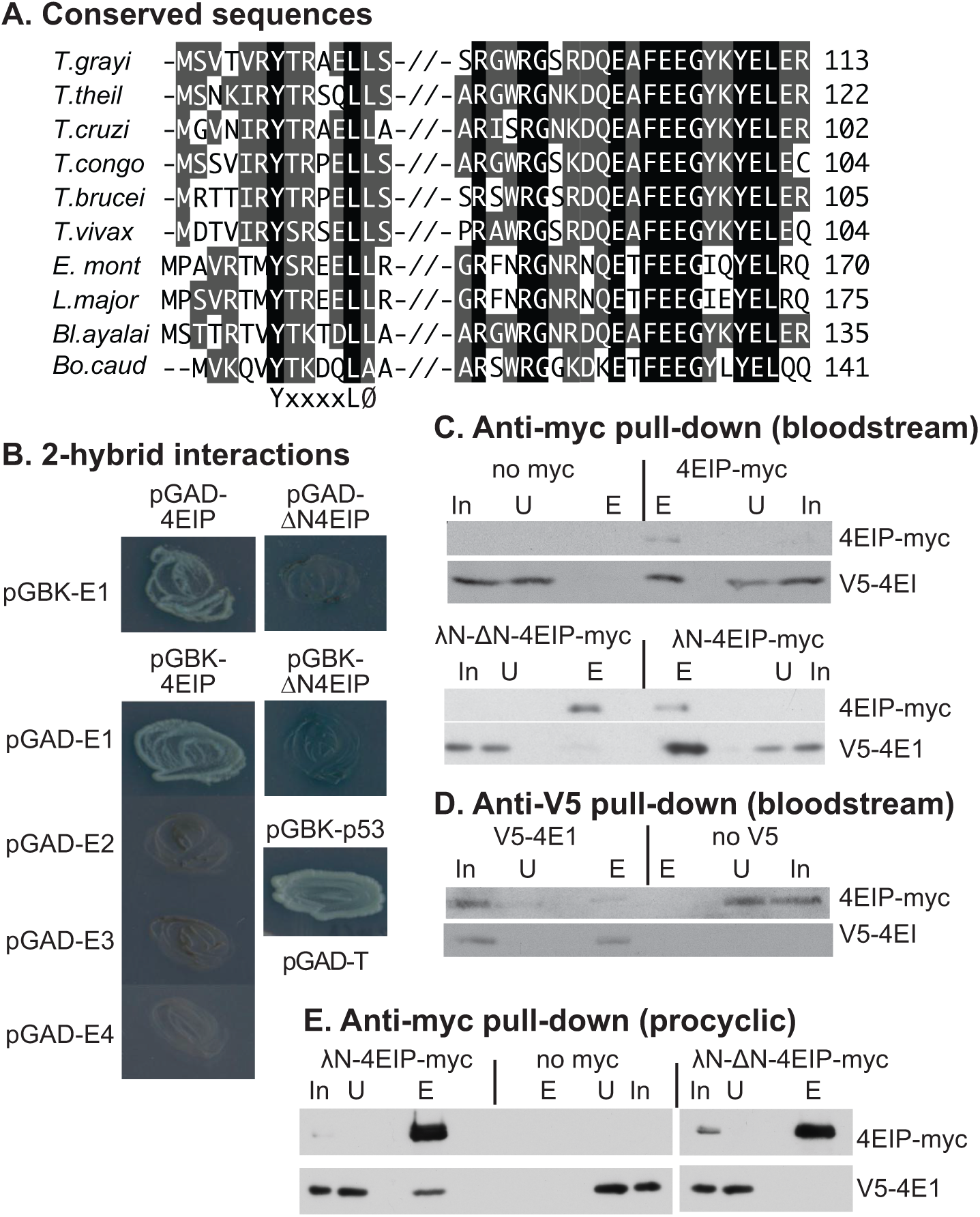
*T. brucei* 4EIP interacts with eIF4E1. A) Conserved regions in kinetoplastid 4EIPs. Amino acids highlighted in black are identical in all aligned sequences. Residues highlighted in grey are partially conserved - either identical or chemically similar in a majority of sequences. Sequences are (in vertical order): *Trypanosoma grayi* DQ04_01981000; *Trypanosoma theileri* ORC92063.1; *Trypanosoma cruzi* TcCLB.508461.290; *Trypanosoma congolense* TcIL3000_9_4530 *Tryanosoma brucei* Tb927.9.11050; *Trypanosoma vivax* TvY486_0905070; *Endotrypanum_monterogeii* EMOLV88_350043200; *Leishmania major* LmjF.35.3980; *Blechomonas_ayalai* Baya_047_0070; *Bodo caudatus* CUG36708.1. B) The photos show growth of yeast on plates with stringent selection for the interaction between bait and prey plasmids. The identities of the plasmids used are shown next to the streaked yeast. Full results with controls are shown in Supplementary Figure S1A. C) Extracts were made from bloodstream-form trypanosomes with or without a V5- *in situ* tagged eIF4E1 and myc-tagged 4EIP. Anti-myc immunoprecipitates were subjected to SDS-PAGE and Western blotting. In: input extract from 5 × 10^6^ cells; U: unbound fraction from 5 × 10^6^ cells; E: eluate from immunoprecipitating beads, from 1 × 10^8^ cells. Upper panel: anti-myc pull-down, with cells lacking a myc-tagged protein as control. Lower panel: anti-V5 pull-down, with cells lacking V5-tagged protein as the control. D) As in (C) but with anti-V5 as the precipitating antibody. E) As in (C) but with procyclic-form trypanosomes.

### *T. brucei* 4EIP interacts with eIF4E1

*L. major* 4EIP was identified as a specific interaction partner of *L. major* EIF4E1 (63). In a 2-hybrid assay, *T. brucei* 4EIP also, as expected, interacted with *T. brucei* EIF4E1, but not with EIF4E2, EIF4E3, or EIF4E4 (Fig 1B and Supplementary Fig S2A). To investigate the interaction inside trypanosomes, we made bloodstream-form *T. brucei* in which one *EIF4E1* open reading frame was N-terminally V5-tagged (V5-4E1). In the same cells, we expressed 4EIP with a C-terminal myc tag (4EIP-myc) (Fig 2B); since 4EIP-myc was expressed from an RNA polymerase I promoter it was probably over-expressed relative to V5-4E1. Anti-myc beads pulled down V5-4E1 from extracts expressing 4EIP-myc, but not from extracts from cells that did not express myc-tagged protein (Fig 1C, upper panel). Similarly, the pull-down of 4EIP-myc by anti-V5 beads was dependent on co-expression of V5-4E1 (Fig 1D).

**Figure 2.**
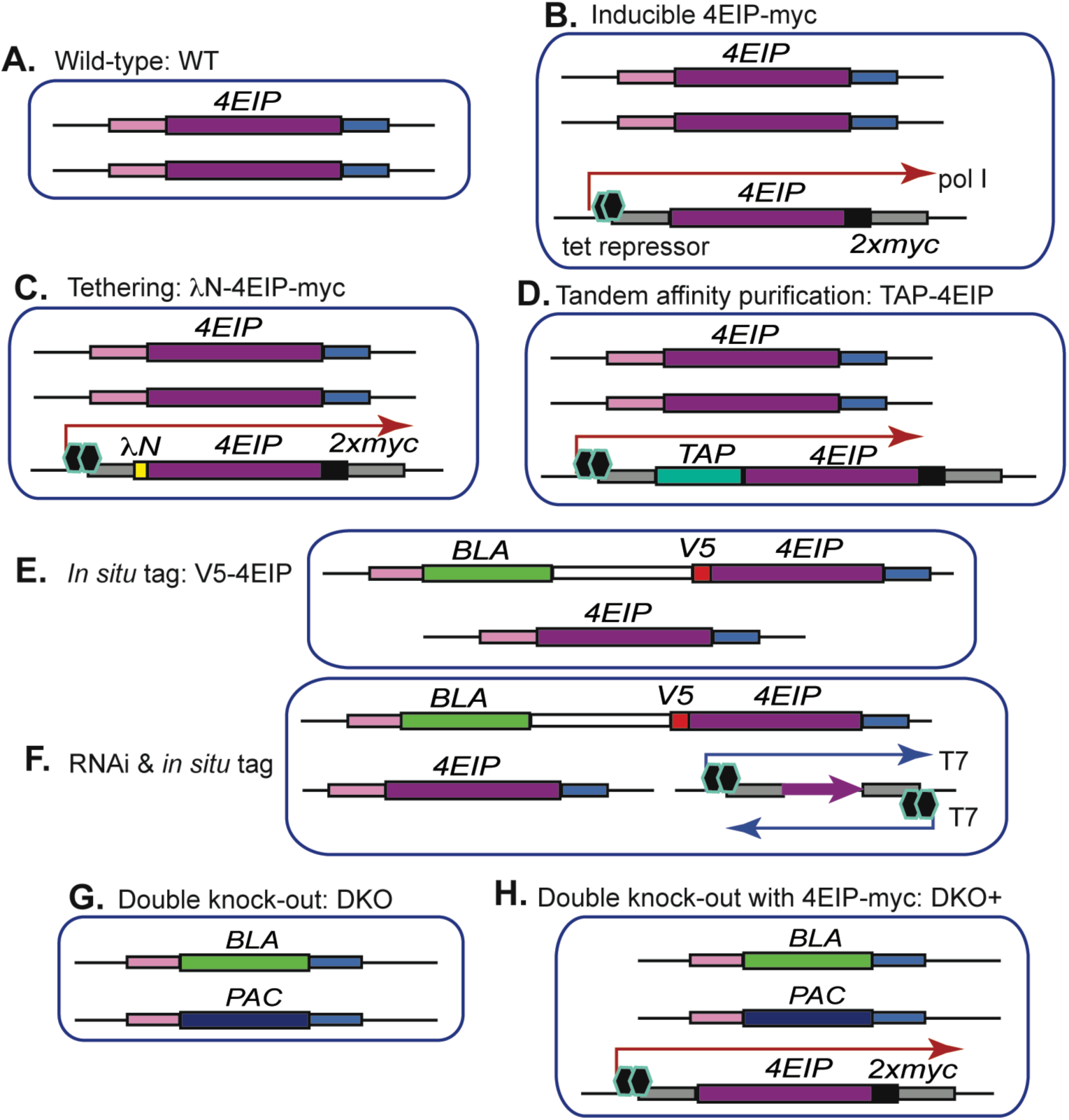
Cell lines. Diagrams are not to scale. Each picture represents the relevant gene loci. The different DNA segments are labelled on the figure.

An 18mer *L. major* peptide that included the YxxxxLØ motif was shown by NMR to interact with *Leishmania* EIF4E1 (63). We therefore tested whether the equivalent sequence in *T. brucei* 4EIP was required for the interaction with *T. brucei* eIF4E1. Indeed, the interaction was no longer seen after deletion of the first 13 amino acids, either by a 2-hybrid assay (Fig 1B, Supplementary Fig S2A, ∆N4EIP) or *in vivo* (Fig 1C). Similar results were obtained in procyclic-form trypanosomes: The EIF4E1-4EIP interaction was present and dependent on the N-terminal peptide (Fig 1E). It is however impossible to rule out the possibility that inside cells, N-terminally truncated but RNA-bound 4EIP is able to interact with cap-bound EIF4E1 in a more transient fashion via additional binding surfaces, with some stabilisation from the linking RNA.

### Suppression of expression by 4EIP does not require the N-terminal EIF4E1-interacting motif or the conserved acidic domain

We next asked which parts of 4EIP are required to suppress expression in the tethering assay. First, we scrutinised existing data from our previous high-throughput screen. In that screen, we expressed a library of protein fragments bearing the lambdaN peptide in cells expressing a boxB reporter mRNA that encoded a toxic enzyme (67). Results for 4EIP from the screen are plotted in Fig 3A. Briefly, each symbol marks the position of the N-terminus of a lambdaN fusion. If the fusion protein suppressed expression, the magenta symbols show higher read counts than the corresponding cyan symbols. The results (67) suggested that protein fragments including the C-terminal 150 residues of 4EIP were sufficient to suppress reporter expression. This region has 20% glutamine and 18% proline, with a predicted pI of 10.4.

**Figure 3.**
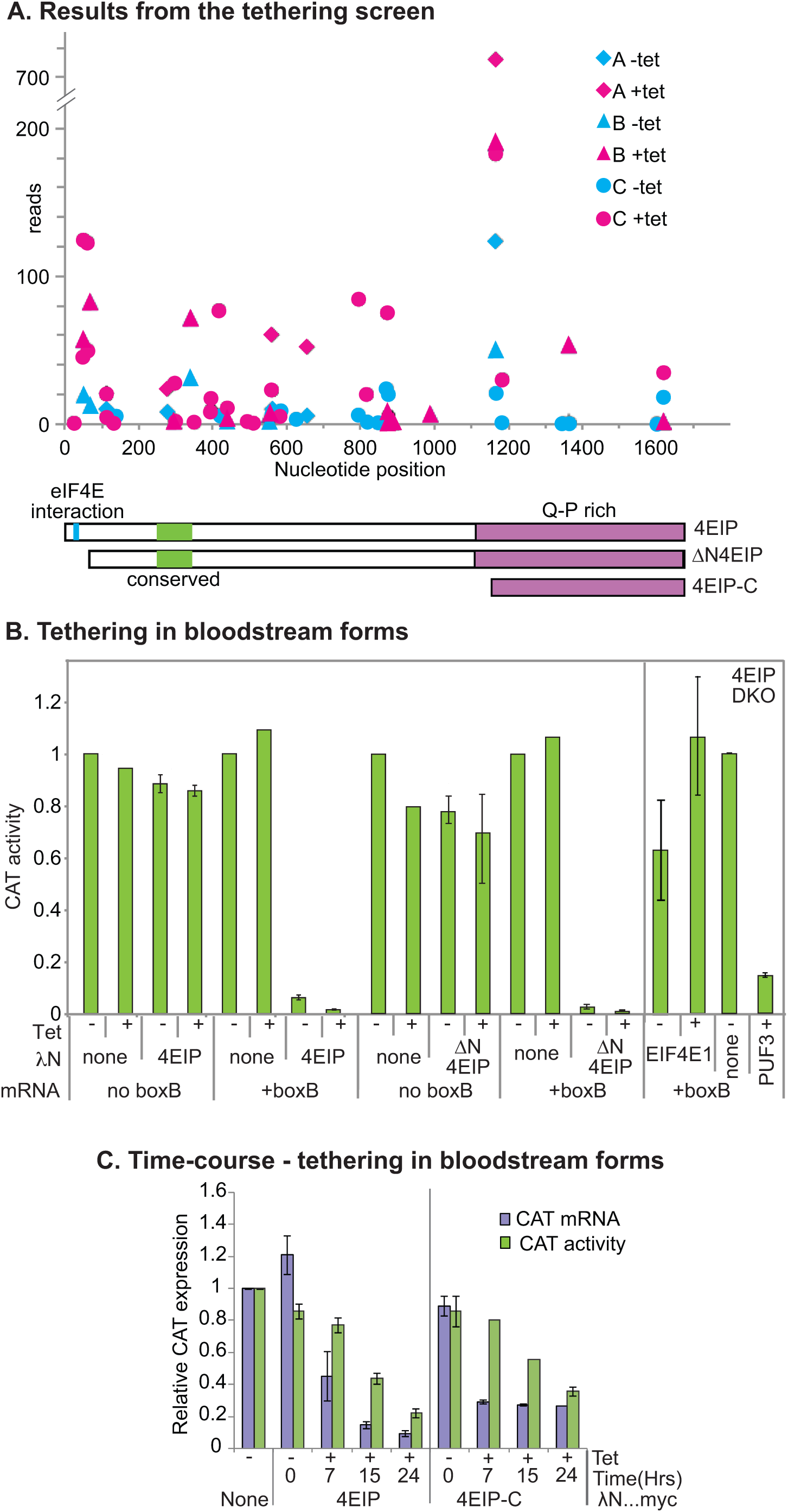
Tethering of 4EIP to an mRNA suppresses expression. A) Results for 4EIP from the published tethering screen (67). The trypanosomes used inducibly expressed an mRNA encoding the glycolytic enzyme PGKB, which is toxic in bloodstream forms (87). They were transfected with a library for inducible expression of lambdaN fusion proteins created from randomly sheared trypanosome DNA. After induction of both *PGKB-boxB* and the lamdaN fusions, cells were grown and the PCR products from the integrated lambdaN fusion proteins were sequenced (67). Results from three independent selections (A, B, C) are shown using different symbols. Counts that are higher in the presence of the inducer, tetracycline, (magenta symbols) than without tetracycline (cyan symbols) indicate suppression of *PGKB* expression. Cartoons of 4EIP showing the two conserved regions (cyan and green) and the PQ-rich C-terminal region are below the x-axis. B) Tethering assay using chloramphenicol acetyltransferase (CAT) reporter. Bloodstream-form trypanosomes expressing a *CAT* mRNA with 5 copies of boxB in the 3’-UTR were used; cells in which the *CAT* mRNA lacked boxB served as control. In the left-had panel, cells were transfected with plasmids designed for tetracycline-inducible expression of either LambdaN-4EIP-myc or LambdaN-4EIP-myc with an N-terminal deletion that eliminated the interaction with eIF4E1 (∆N4EIP). CAT activity was measured in the presence and absence of tetracycline. Results are expressed as arithmetic mean ± standard deviation of at least 3 measurements. Expression of myc-tagged protein is shown in Supplementary Fig S3A. Right-hand panel - cells lacking 4EIP were used to test the activity of tethered EIF4E1. Expression of the EIF4E1, along with a typical Northern blot (done twice) is shown in Supplementary Fig S3B. C) Time course of the tethering effect in bloodstream forms, using either 4EIP or the C-terminal domain.

To confirm the screening results we used bloodstream forms that contained a constituteively transcribed *CAT-boxB* mRNA. We first expressed full-length lambdaN-4EIP-myc from a tetracycline-inducible promoter (Fig 2C, Supplementary Fig S3A). Even in the absence of tetracycline, *CAT* mRNA was not detectable and CAT activity was suppressed by about 95% (Fig 3B). The suppression without tetracycline can be explained by leaky lambdaN-4EIP-myc expression (Supplementary Fig S3A, arrow). The same result was obtained upon tethering of ∆N4EIP (Fig 3B, Supplementary Fig S3A). For both proteins, expression of a *CAT* mRNA without boxB was unaffected (Fig 3B). Thus tethered 4EIP acts independently of the EIF4E1 interaction motif, as was previously seen in Opisthokonts.

We then, using new lines, examined the time course of the tethering effect, comparing the full-length protein with the C-terminal 150 residues (Fig 3A). The kinetics of RNA decrease were similar for the full-length protein and for the C-terminal domain (Fig 3C). CAT activity declined more slowly than RNA as expected, because we are here also observing degradation of CAT protein that was synthesised prior to tetracycline addition (Fig 3C). These results confirmed that as predicted from the screen, the C-terminal domain of 4EIP is sufficient for suppression of gene expression by tethered 4EIP.

Results so far indicated that the activity of 4EIP in the tethering assay was independent of the EIF4E1 interaction. We therefore asked the complementary question - whether the action of EIF4E1 depended on 4EIP. EIF4E1 had highly suppressive activity in the tethering assay, both in high-throughput screens and when tested individually (67). To find out whether this activity required 4EIP we repeated the tethering assays in cells lacking 4EIP. (These “DKO” cells are described later in this paper.) A control tethered protein, PUF3, was able to suppress reporter expression in ∆4EIP cells (Fig 3B, far right). The expression of LambdaN-EIF4E1 was poorly regulated by tetracycline (2-5-fold) (Supplementary Fig S3B) but interestingly, in all three replicates CAT activity was higher in the presence of tetracycline than in its absence (Fig 3B, 1.2 - 2.3-fold) but not significantly different from the control cell line lacking tethered protein. In the absence of 4EIP, therefore, tethered EIF4E1 is unable to repress translation. These results are exactly the same as those previously seen for mammalian 4EHP and GIGYF2: the suppressive activity of tethered 4EHP is absent in mutant mammalian cells that lack GIGYF2, but tethered GIGYF2 represses in the absence of 4EHP binding (24).

### 4EIP tethering inhibits translation initiation and causes mRNA decay

We next repeated the 4EIP tethering experiment in procyclic forms (Supplementary Fig S3C). Expression of lambdaN-4EIP-myc in procyclic forms resulted in mild growth inhibition (Fig 4A). Induction of lambdaN-4EIP-myc expression for 24h resulted in a ten-fold decrease in *CAT-boxB* mRNA, and a more moderate decrease in CAT activity: CAT protein probably persisted from the pre-induction period. LambdaN-GFP-myc had no effect, while tethering of poly(A) binding protein 1 (PABP1) increased expression, as expected (70) (Fig 4B)

**Figure 4.**
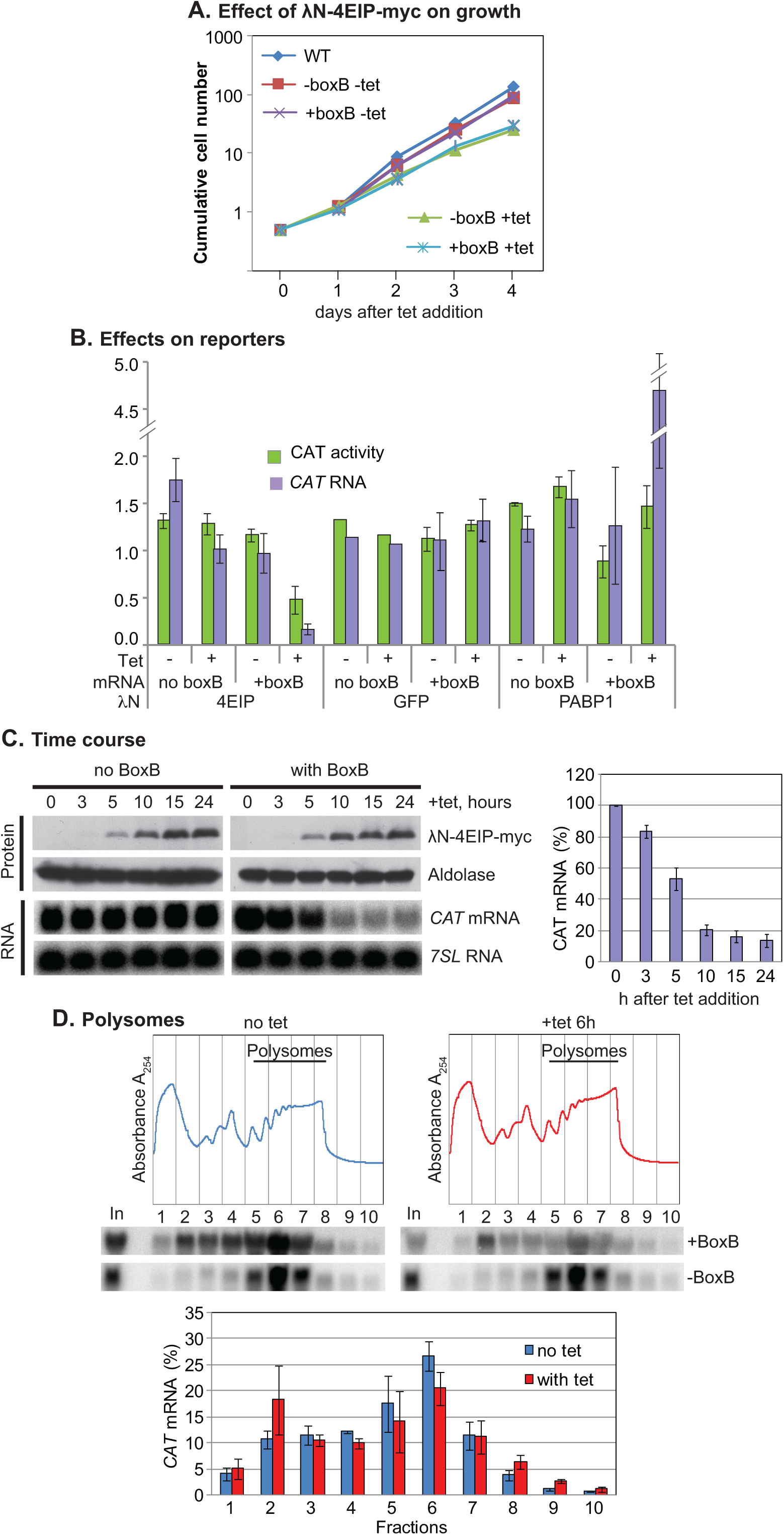
4EIP tethering suppresses translation and induces mRNA destruction. A) Effect of LambdaN-4EIP-myc expression on cell proliferation. A cumulative growth curve is shown. Results are arithmetic mean with error bars showing standard deviation from three assays. B) Effects of different tethered reporters on CAT activity (green) and mRNA (mauve). Results are arithmetic mean with error bars showing standard deviation. Fusion protein expression is shown in Supplementary Fig S3C. C) Time course of LambdaN-4EIP-myc expression in cells expressing *CAT* mRNA with or without boxB. A Northern blot showing the effect on *CAT* mRNA is shown below, and quantitation of replicates is to the right. D) Polysomes were separated on sucrose gradients, either without tetracycline (blue), or 6h after tetracycline addition (red). Typical gradient profiles and Northern blots are shown at the top, and quantitation of three replicates beneath.

To find out whether 4EIP inhibits translation, we needed to find a time point at which expression of LambdaN-4EIP-myc was evident, but *CAT-boxB* mRNA was still detectable. This occurred between 5 and 10h after induction (Fig 4C). Analysis of sucrose gradient fractions revealed that even without induction, much more of the *CAT-boxB* mRNA was in the low density fractions than was the case for the mRNA without boxB. After 6h induction, the boxB-containing mRNA shifted further towards the lighter part of the gradient, with the amount of *CAT* mRNA in the polysomal fraction showing a preferential decrease (Fig 4D). There are two possible interpretations of this result. One is that attachment of 4EIP to the reporter mRNA represses its translation, with subsequent degradation. The alternative possibility is that 4EIP stimulates deadenylation or decapping with a decrease in translation as a secondary effect. There was, however, no evidence for mRNA shortening from the Northern blots (Fig 4C) and we would expect de-capped mRNA to be destroyed very rapidly.

### 4EIP interactions with other proteins

In order to find out how 4EIP causes mRNA decay or translation repression we looked for interaction partners beyond EIF4E1. We used bloodstream-form trypanosomes inducibly expressing 4EIP with an N-terminal tag suitable for tandem affinity purification (TAP) (Fig 2D, Supplementary Fig S4), in triplicate and with four GFP-TAP controls. Some experiments were also done using 4EIP-myc. Purifications were done either in the presence of an RNase inhibitor, or with RNase. As expected, eIF4E1 consistently co-purified with 4EIP (Supplementary Table S1). The remaining co-purifying proteins were mostly very abundant and are regularly found as contaminants in our mass spectrometry analyses. Some of them are also unlikely to be associated with 4EIP based on their intracellular location. No promising novel interactions were found that might have explained the 4EIP mechanism. It is interesting that although it was excluded from our final list by our selection criteria, EIF4A1 was enriched in 5 out of 6 4EIP-TAP preparations.

Since mass spectrometry routinely detects abundant proteins, and results are dependent on the life-cycle stage, we also screened a random-shotgun yeast 2-hybrid prey library (53) with 4EIP bait, to identify possible direct interaction partners. Results were assessed initially by sequencing individual clones, then by sequencing of pooled library inserts (Supplementary Table S2, Supplementary Fig S2B). We compared the results for 4EIP with those from four components of the CAF1-NOT complex (Supplementary Table S2) (71). For EIF4E1 the only interacting fragment began at the start codon (Supplementary Table S2). There were 32 possible new 4EIP interaction partners, defined as genes with two or more interacting fragments that interacted either with one or none of the other four baits. Among these, proteins with links to RNA metabolism were the RNA-binding domain protein ZC3H14, an RNaseH domain protein, the spliceosome subunit U5-40K, polyadenylation complex protein CstF50, and Dicer 1. In the inital tethering screen, using protein framgents, none of these five proteins could suppress expression of the reporter (67). In contrast, full-length tethered ZC3H14 was indeed capable of suppressing reporter expression (68). It was not, however, detected in a stringent purification of proteins that could be cross-linked to mRNA (mRNP proteome) (68) and was not detected using 4EIP-TAP purification.

We tested three candidate interactors from the yeast 2-hybrid screen by co-immunoprecipitation: EF2 (apparently specific), RBP9 and NOT1, which each had only one interacting fragment but can suppress when tethered. Interactions were tested with either 4EIP-myc (Fig 2B) or 4EIP with an N-terminal V5 tag (Fig 2E). None of the results was positive (Supplementary Figure S5). When two fragments covering NOT1 were used as bait, 4EIP was also not detected (68). In total, none of the experiments provided any evidence for association of 4EIP with known components of the mRNA degradation machinery.

### mRNAs associated with 4EIP are relatively unstable

To find out which mRNAs are associated with 4EIP, we performed two independent single-step purifications of TAP-tagged 4EIP (Supplementary Fig S4B). We sequenced the 4EIP-associated RNAs and compared them with those in the unbound fraction (Supplementary Table S3, sheet 2). We found 219 mRNAs that were at least 2-fold enriched (as judged from normalised read ratios) in both pull-downs. We compared these with 399 mRNAs that were reproducibly less abundant in the bound than in the unbound fraction (Supplementary Table S3, sheet 1). Results are shown in Figure 5A-C. The mRNAs encoding ribosomal proteins (which were not bound, see below and Supplementary Table S3, sheet 1) are plotted separately because they behave differently from most other mRNAs in that they are poorly translated but extremely stable, and unlike other mRNAs, always excluded from stress granules (72). The 4EIP-bound mRNAs had slightly lower ribosome densities than the unbound ones (Fig 5A), but more dramatically, their median half-life was half that of the unbound mRNAs (Fig 5B). These results are consistent with a suppressive role for 4EIP. Strangely, though, the 4EIP-bound mRNAs had coding regions that were on average, less than half the length of those of unbound mRNAs (Fig 5C). We therefore worried that the relative instabilities of bound mRNAs were an artifact that was in some way linked to short coding region length. To check this, we re-analyzed a size-matched subset of bound and unbound mRNAs (Fig 5F; Supplementary Table S3, sheet 4). The differences in ribosome density and half-life persisted, showing that this is an intrinsic property of 4EIP-bound mRNAs (Fig 5D, E). No functional protein category was enriched in the bound fraction. There was also no correlation with developmental regulation. The RNA-binding protein RBP10 causes degradation of mRNAs in bloodstream forms (73). Among the 232 bound mRNAs was that encoding EP procyclin, along with 10 other targets of RBP10; however the “unbound” set also included 9 RBP10 targets. The annotated 5–-untranslated region lengths of bound and unbound mRNAs were similar (medians of 84 and 94 nt respectively) and using DREME, no enriched sequence motifs were found in either the 5’ or 3’-untranslated regions of the bound mRNAs.

**Figure 5.**
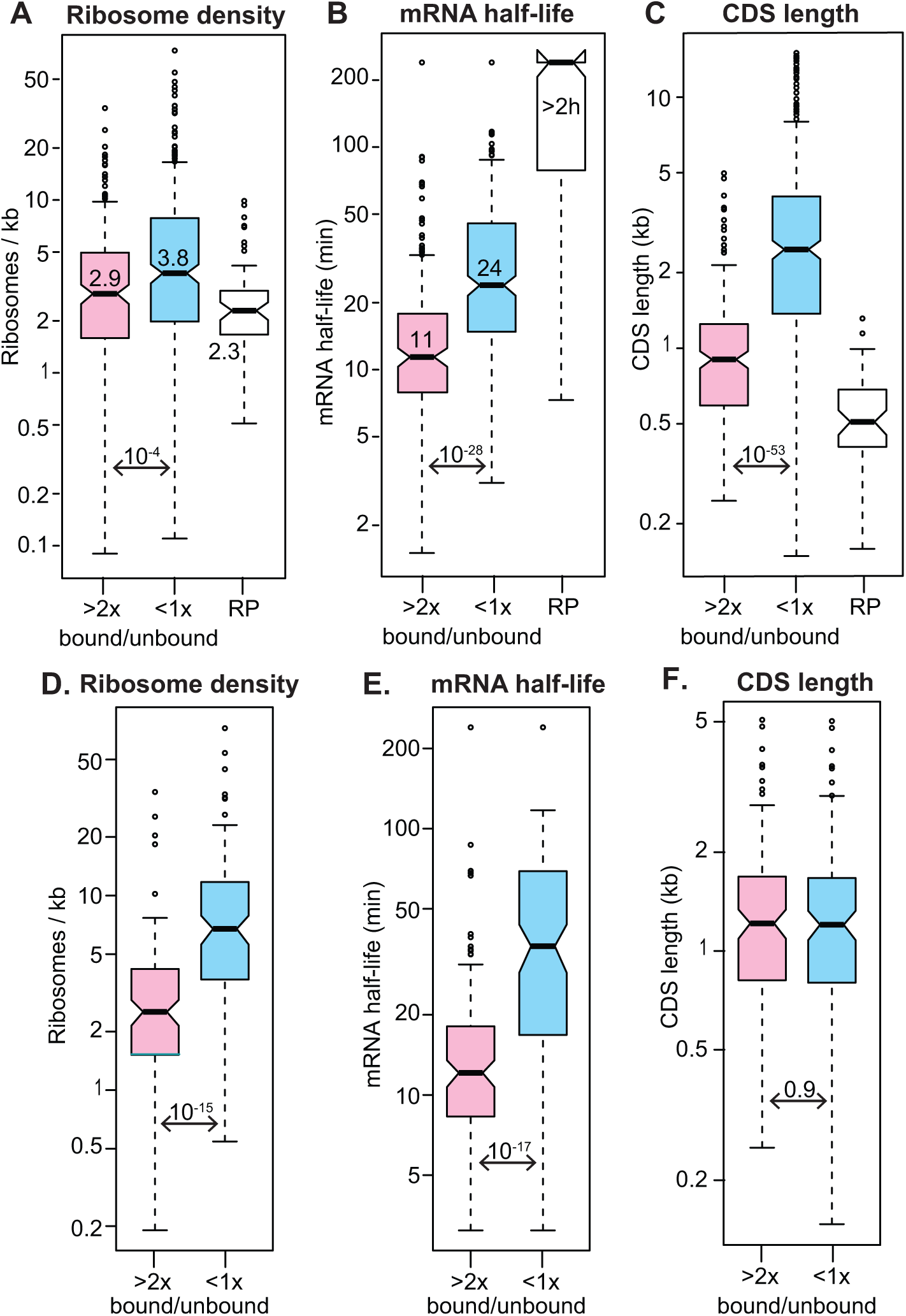
RNAs bound to 4EIP are relatively unstable. mRNAs were classified as "bound" if the reads per million in the fraction bound to TAP-4EIP were at least twice those in the flow-though in both biological replicates, and the minimum count number was 10. "Unbound" mRNAs has a ration of less than 1 in both replicates. The mRNAs encoding ribosomal proteins (RP) were mostly not bound but were considered separately. All results are for bloodstream forms, A) Ribosome density for all "bound" and "unbound" mRNAs, as judged by ribosome profiling (49). Medians are indicated, as are the student t-test values. B) mRNA half-lives for all "bound" and "unbound" mRNAs (47) C) Open reading frame or coding sequence (CDS) lengths of all “bound” and “unbound” mRNAs. D) As (A), but for a subset of length-matched mRNAs (Supplementary Table S3 sheet 4) E) As (B), for the subset of length-matched mRNAs F) As (C), for the subset of length-matched mRNAs

More than half of all ribosomal protein mRNAs were in the unbound fraction (Bonferroni-corrected Fisher test P value 10^−30^) and only one was in the bound fraction. Their lack of association with 4EIP shows that their poor translation is not caused by 4EIP. The only other functional groups that were significantly enriched in the unbound fraction were glucose and glycerol metabolism (mostly glycolytic enzymes, corrected P-value 0.02) and lysosomal proteins (corrected P-value 0.05). In addition, the "unbound" list included mRNAs encoding 4 of the 8 TRiC complex components, and, among the cytoskeletal proteins, many flagellar structure components (including tubulin) but only 3 motor proteins.

We also statistically analysed the results using DeSeqUI (74), an R package that runs DEseq2 (75) on trypanosome data (Supplementary Table S3, sheet 2 and Supplementary Fig S6). Although this programme is not designed for pull-down experiments, the results were generally similar to those calculated from read ratios. The 232 “bound” mRNAs (Supplementary Fig S6A) included 177 of those that had been found previously, with a slight bias towards shorter mRNAs (Supplementary Fig S6B). The bias towards short open reading frames in the bound fraction was probably not caused by short open reading frames being more poorly translated: in fact overall, longer open reading frames have lower ribosome densities than shorter ones (Supplementary Fig S6C). Ribosomal protein mRNAs were again significantly enriched in the unbound category (Supplementary Fig S6D).

### 4EIP is not essential in bloodstream-form trypanosomes

To investigate the role of 4EIP in trypanosomes, we first targeted it by RNAi in monomorphic bloodstream forms that expressed *in situ* V5-tagged 4EIP (Fig 2F). RNAi was effective, although a little tagged protein persisted (Fig 6A and Supplementary Fig S7A). There was a small but reproducible increase in the division time (Fig 6A). For unknown reasons, we were unable to obtain procyclic form clones with effective RNAi.

**Figure 6.**
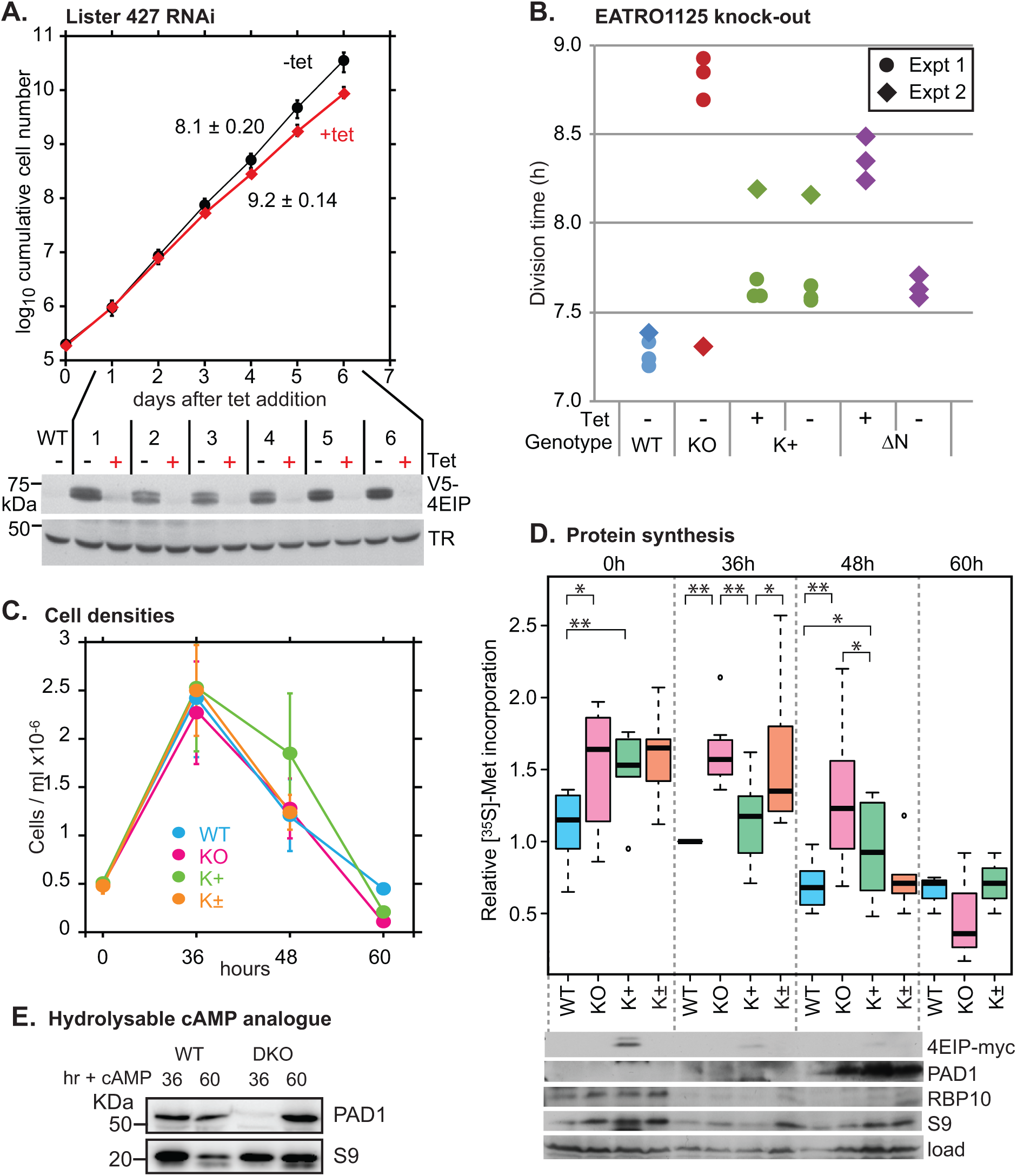
Cells without 4EIP differentiate poorly at high density. A) Depletion of 4EIP in monomorphic bloodstream forms (Lister 427) results in a very mild growth defect. Cumulative growth of cultures with tetracycline-inducible RNAi is shown, with mean and standard deviation for three different cell lines. Expression of V5-4EIP in one of the three different RNAi lines, analysed by Western blotting, is shown below. TR is trypanothione reductase. Results for the other two cell lines are in Supplementary Fig S7A. B) Division times of different EATRO1125 cell lines. The division times of the same cell lines were measured in two independent experiments. Results for the individual replicates are shown. KO: no 4EIP; K+: knock-out line with inducible 4EIP-myc, with or without tetracycline; ∆N - knock-out line with inducible lambdaN-∆N-4EIP-myc, with or without tetracycline; e. Tetracycline was added 48h prior to the start of measurements. In the second experiment (diamonds) the knock-out line appeared to have adapted to the lack of 4EIP during continuous culture. C) Growth of different EATRO1125 cell lines allowed to reach high density. Cells with a starting density of 5×10^5^/ml were allowed to grow in HMI-9 media containing 1.1% methylcellulose for 60 hrs. Results are mean and standard deviation of three replicates. K+ are the complemented cells K± are the K+ cells grown without tetracycline. D) [^35^S]-Methionine incorporation into proteins during differentiation of EATRO1125 at high density. Trypanosomes were cultured as shown in (B), washed and pre-incubated for 1 hr in labelling medium at 37°C. Methionine (^35^S) was then added for 20 min and proteins were analyzed by SDS-PAGE (Supplementary Figure S8B). Results for seven independent experiments are shown; four of the experiments included time=0h and three included time = 60h. The colour code is a paler version of (B). Student t-test results are: ** p<0.01, * p<0.05, both as paired or unpaired tests. Expression of various proteins, by Western blotting, is shown under the box plot. E) EATRO1125 Cells lacking 4EIP show a delayed response to a cell-permeable cAMP analogue. 8-pCPT-2’-O-Me-5’-cAMP was added to growing cells at 9247 cl:16210^5^ /ml to a final concentration of 10 μM and Western blots were made after 36h and 60h.

To find out whether 4EIP was essential for survival, we deleted the genes in both Lister 427 and pleomorphic EATRO1125 bloodstream forms. The deletion was successful in both cases; evidence is shown in Supplementary Fig S7C and D. For Lister 427 cells, the division time increased from 8h (wild-type and single knock-out) to 10.4h. For the EATRO1125 ∆*4eip*/∆*4eip* knockout cells, a mild growth defect was observed (Fig 6B, “KO”). A single repeat measurement after several months of continuous culture suggested, however, that the cells had been able to adapt to life without 4EIP (Fig 6B, diamond symbols). We introduced a tetracycline-inducible 4EIP-myc gene into the EATRO1125 ∆*4eip*/∆*4eip* knockout cells (Fig 2H, Supplementary Fig S7E) and this restored normal growth even in the absence of tetracycline (Fig 6B, K+). We do not know how much 4EIP-myc is expressed relative to the endogenous amount. However, we do know that RNA polymerase I transcription is 5-10 times more active than transcription by RNA polymerase II (76). Since the level of 4EIP-myc in the absence of tetracycline was 5±3% of the induced level, (Supplementary Fig S7E), it is quite possible that this “uninduced” amount was sufficient for complementation. N-terminally GFP-tagged 4EIP is located in cytosolic granules in transiently starved procyclic forms (see http://tryptag.org/?query=Tb927.9.11050) (77). 4EIP-myc had a similar location in the complemented bloodstream and procyclic forms (supplementary Fig S7F).

In cells with inducible expression of 4EIP with the N-terminal 11-amino-acid deletion (∆N), low-level expression in the absence of tetracycline (Supplementary Fig S7E) was sufficient to complement the growth defect while induced expression gave slower growth (Fig 6B), suggesting that excess expression of the deleted version was toxic.

### 4EIP is required for normal translation arrest during stumpy form differentiation

We now attempted to induce stumpy-form differentiation in the EATRO1125 cells. The protocols used were as previously described (73). Cultures in methyl cellulose were started at 5 ×10^5^/ml and allowed to grow for 48-60h. Cell numbers peaked within 36h then started to decline (Fig 6C, Supplementary Fig S8A). The complemented knock-out cells, which were probably over-expressing 4EIP, consistently grew to slightly higher maximum densities and survived for longer before numbers started to decline (Fig 6C. Supplementary Fig S8B). After 36h, we examined the cells by phase-contrast and fluorescence microscopy with DNA stain. All cultures looked similar. They consisted predominantly of G1-arrested cells; roughly 2% of cells were in the S or G2 phases of the cell cycle and variable numbers of abnormal-looking cells were also present. All 48h cultures expressed the stumpy-form marker PAD1 (Fig 6D, Supplementary Fig S8D; for a control negative results see Fig 7) but expression in the knock-out was often lower than in the other lines (Fig 7A). There was no significant difference between the knock-out and the complemented trypanosomes in the distance between the nucleus and kinetoplast (Supplementary Fig S8C). All of the cultures lost expression or RBP10, a bloodstream-form regulator, within 36h (Fig 6D).

**Figure 7.**
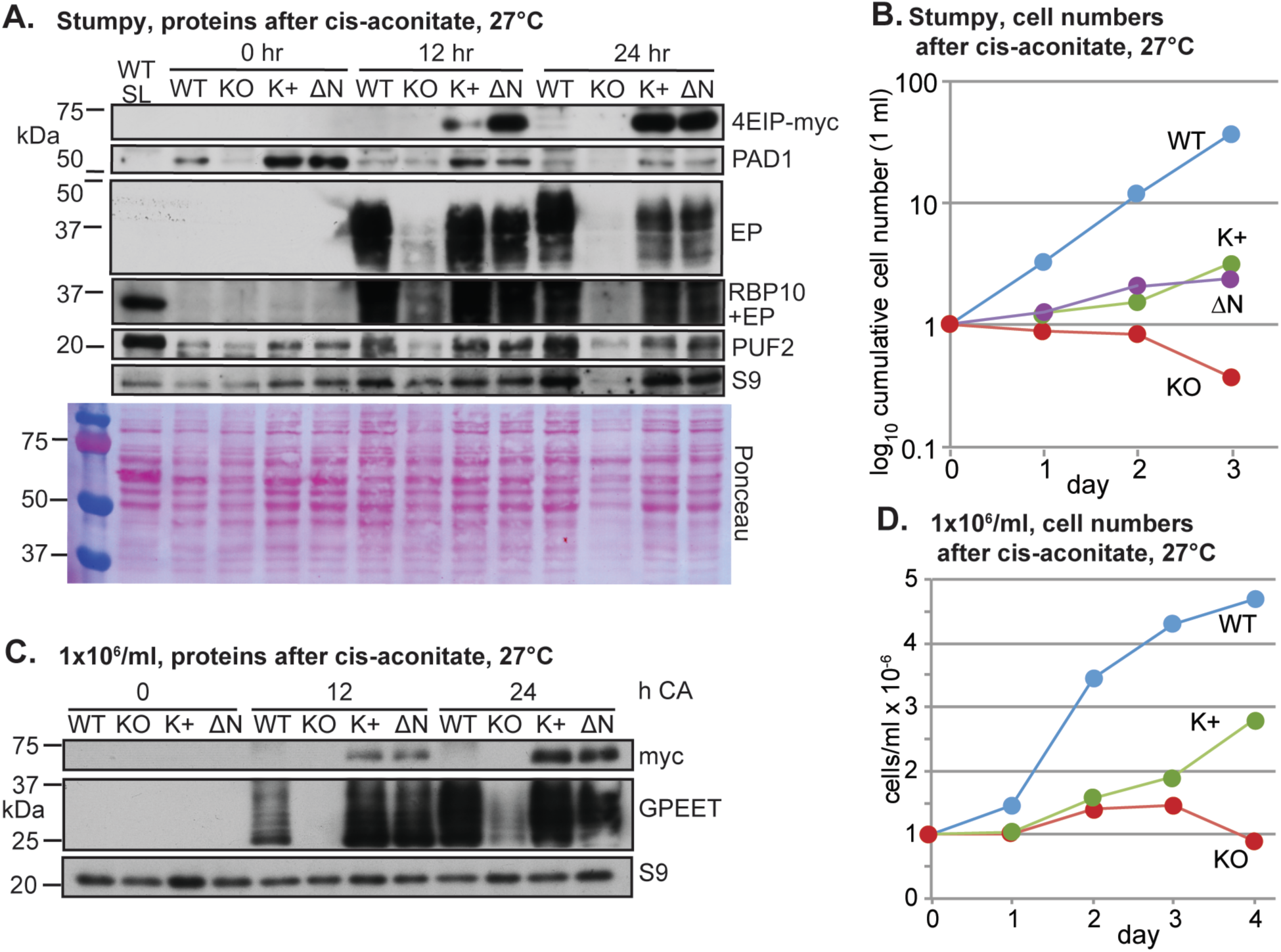
Cells without 4EIP differentiate poorly into procyclic forms. A) Expression of various proteins in EATRO1125 cells cultured to high density for 60h, then treated with cis-aconitate (CA) and transferred to 27°C. WT SL is long slender bloodstream forms; KO: no 4EIP; K+: knock-out line with inducible 4EIP-myc and tetracycline; ∆N - knock-out line with inducible lambdaN-∆N-4EIP-myc, with tetracyline. B) Cell counts for cells treated as in (A) (to the end of day 1), then transferred to procyclic medium C) Expression of various proteins in cells cultured to 1 × 10^6^/ml, then treated with cis-aconitate and transferred to 27°C. Replicates are in Supplementary Fig S9C. D) Cell counts for cells from C, transferred to procyclic medium on day 1.

Stumpy formation is accompanied by a general decrease in protein synthesis (34). To find out whether the lack of 4EIP affected this, we measured incorporation of [^35^S]-methionine into proteins. After 36h the knock-out cells consistently showed higher protein synthesis than either the wild-type or the complemented controls (Fig 6D); but the difference was less marked after 48h and had disappeared at 60h. Examination of the protein profiles did not reveal any highly synthesized proteins that were consistently different between the cell lines (Examples are in supplementary Fig S8D).

Stumpy-form differentiation in the above experiments was caused by density-dependent signaling, but it can also be induced by cell-permeable, hydrolysable cAMP analogues (78). To test this, 8-pCPT-2’-O-Me-5’-cAMP was added to growing cultures. Cells lacking 4EIP showed a delayed response, as measured by PAD1 expression (Fig 6E).

From these experiments we concluded that loss of 4EIP results in a delay in the inhibition of gene expression that accompanies stumpy formation. We attempted to compare mRNAs from the different time points but were unfortunately unable to obtain intact RNA from the 36h and 48h cultures. The most likely explanation is that the cell pellets contained very high levels of RNase, either from dead cells or from live stumpy forms themselves. Nevertheless, from the other results it seems likely that the major role of 4EIP in stumpy formation is in suppressing translation.

### 4EIP is required for normal differentiation to procyclic forms

Next, we investigated differentiation of the bloodstream forms into procyclic forms. 48h after the cells attained a density of 1 x 10^6^/ml, cis-aconitate (final concentration 6mM) was added and the temperature was reduced to 27°C. At this point (time = 0 in Fig 7), the myc-tagged complementing proteins were not detected, presumably because stumpy-form cells have extremely low transcription and translation (33). However, within 12h of cis-aconitate addition, the complementing proteins were again seen. After 12h cis-aconitate, expression of the procyclic surface protein EP procyclin was clearly visible in wild-type cells, but barely detectable in the ∆*4EIP* knock-out (Fig 7A, supplementary Fig S9A), and the same was true after 24h. Expression of either 4EIP-myc or ∆N4EIP-myc in the ∆*4EIP* cells restored EP procyclin expression. Expression of RBP10, a bloodstream-form-specific RNA-binding protein, was lost within 24h (supplementary Fig S9A). After 24h in cis-aconitate, the cells were transferred to procyclic medium at 27°C. The wild-type cells grew, as previously described (73). In contrast, the ∆*4EIP* knock-out cells lingered in the cultures for up to a week but were unable to recover. The complemented cells had intermediate behaviour, showing that at least some of the differentiation defect in the ∆*4EIP* knock-out cells was due to the lack of 4EIP (Fig 7B). The difference between the complemented cells and wild type is most likely due to loss of other aspects of differentiation competence during the three rounds of clonal selection (see also Supplementary Fig S9B).

Results so far indicated that 4EIP is required for formation of stumpy forms that are competent for full procyclic-form differentiation. We therefore wondered whether the cells would be able to differentiate if they had not been left at high density for a prolonged period. To test this, the trypanosomes were grown to 1 ×10^6^/ml, then cis-aconitate was added and the temperature was shifted to 27°C. Once again, a delay in EP or GPEET procyclin expression was specifically observed in the ∆*4EIP* knock-out cells (Fig 7C, Supplementary Fig S9B). After 24h the medium was changed to procyclic form medium. The knock-out cells were again unable to resume replication (Fig 7D), in contrast to the complemented cells. The 8-pCPT-2’-O-Me-5’-cAMP-treated cells showed the same defect (Supplementary Fig S9D).

These results showed that 4EIP is required for differentiation of bloodstream to procyclic forms. 4EIP is not essential in established procyclic forms (E. Freire, Fiocruz, Brazil, personal communication). The results therefore suggest that the defect is caused by an inability of the cells to suppress expression of a subset of proteins during the actual differentiation process. Notably, the defect could be complemented by N-terminally deleted 4EIP, implying that the essential function of 4EIP during differentiation does not require it to interact with EIF4E1.

### EIF4E1 is not essential

It was previously reported that while procyclic forms with RNAi targeting EIF4E1 showed only mildly slower growth (55), bloodstream-form trypanosomes with depleted EIF4E1 were unable to grow (55). Essentiality of EIF4E1 in bloodstream forms would suggest that it has a function that is independent of 4EIP. We, however, observed only a slight growth defect after RNAi (Fig 8A). Moreover, using differentiation-competent EATRO1125 bloodstream forms, both *EIF4E1* gene copies were readily deleted (Supplementary Fig S10A, B), with only a mild growth defect (Fig 8B). We do not understand why the published RNAi was so deleterious: although the dsRNA that was used comprises the complete open reading frame, it has no predicted off-target matches.

**Figure 8.**
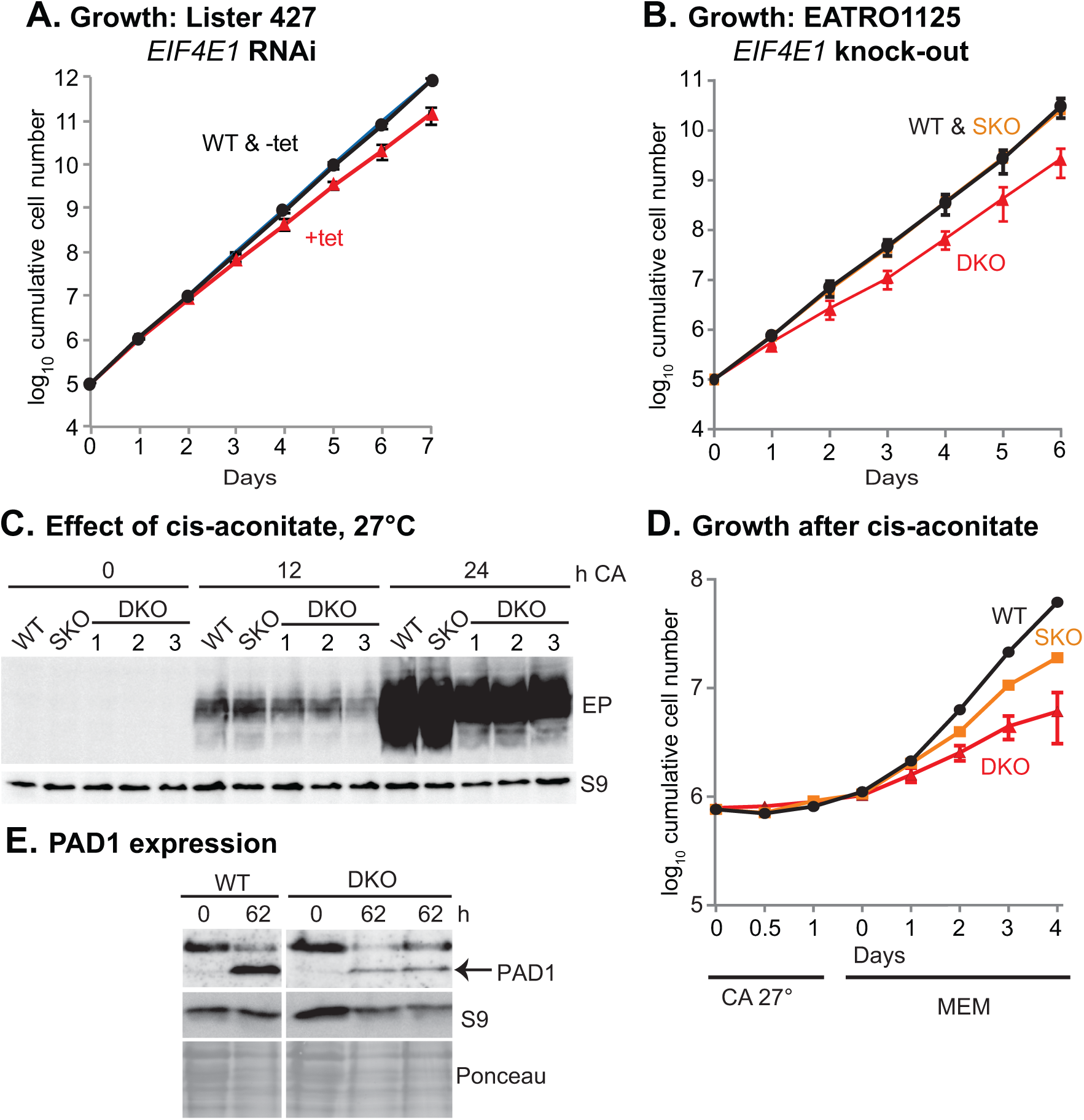
Cells without EIF4E1 have a mild growth defect. A) Depletion of EIF4E1 in Lister 427 bloodstream forms results in a very mild growth defect. Cumulative growth of cultures with tetracycline-inducible RNAi is shown, with mean and standard deviation for 3 independent experiments. B) Cumulative growth of different EATRO1125 cell lines. For WT and SKO (single replacement), the division times of the same cell lines were measured in two independent experiments. For DKO (both genes replaced), results for 3 different lines are shown. C) Effect of adding cis-aconitate (CA) and shifting the temperature to 27°C (EATRO1125). The starting cell density was 1×10^6^/ml. EP procyclin expression was measured by Western blotting, with ribosomal protein S9 as the loading control. Although it is not evident from the image, the last 3 lanes are somewhat under-loaded, and quantitation revealed no significant differences between DKO lines and the SKO or WT. D) Growth of cells from (C) after transfer to medium suitable for procyclic forms (MEM). E) Expression of PAD1 after 62h incubation at high density. Results at 62h for two different DKO clones are shown. No reproducible differences in PAD1 expression were seen.

Since the function of 4EIP in differentiation appeared to be independent of interaction with EIF4E1, we next tested whether the *EIF4E1* knock-out cells were able to differentiate. After growth to 1 ×10^6^/ml, followed by cis-aconitate addition, the kinetics of EP procyclin expression were similar to wild-type (Fig 8C). Upon transfer to procyclic medium, cells with a single *EIF4E1* gene recovered slower than the wild-type, and cells lacking EIF4E1 were slower still, but they could nevertheless reproduce (Fig 8D). PAD1 expression was also seen after incubation at high density (Fig 8E). These results supported our previous conclusion that EIF4E1 is not required for 4EIP to act during differentiation.

## Discussion

Our results indicate that tethering of *T. brucei* 4EIP to an mRNA suppresses its translation and promotes RNA degradation. In bloodstream forms, 4EIP is preferentially bound to mRNAs with relatively short half-lives and lower ribosome densities, suggesting that natural binding has a similar effect. We do not know which of the two effects on mRNA is the primary activity of 4EIP: they could be linked, since poor translation initiation might result in mRNA decay but deadenylation or decapping can prevent translation. No interactions of 4EIP with the mRNA degradation machinery were detected.

Although 4EIP appears - from UV cross-linking results - to bind mRNA directly, it lacks a known RNA-binding motif and is not clear how the protein is recruited to specific mRNAs: we found no common sequence motifs in the bound transcripts. In mammalian cells, 4EHP can be recruited to target mRNAs via specific RNA-binding proteins (22,23). Only one putative RNA-binding protein, ZC3H14, interacted with 4EIP in a yeast 2-hybrid screen. ZC3H14 can indeed suppress expression in the bloodstream-form tethering assay, but there is as yet no evidence from either ribosome profiling (48,49) or proteomics (79,80) that ZC3H14 protein is expressed. (Might ZC3H14 be specific to life-cycle stages that have not yet been molecularly characterised?) The mechanism(s) by which 4EIP recognises its targets thus remains to be discovered.

The only 4EIP partner protein that emerged reliably from our analyses was EIF4E1. There are various possible mechanisms by which 4EIP and EIF4E1 might act. One possibility is that without 4EIP, EIF4E1 can act as a translation initiation factor (Fig 9A). *L. major* EIF4E1 can interact with EIF3 in an *in vitro* pull-down, and the interaction was also detected *in vivo*, although over-expression of EIF4E1 was required (63,64). An argument against this idea is that when EIF4E1 was tethered in the absence of 4EIP, there was no convincing increase in reporter expression, whereas tethered EIF4E3/G4 and EIF4E4/G3 enhance expression considerably (67,68). However, the presence of the tags, or the fact that the protein was tethered to the 3’-UTR, could have affected EIF4E1 function. *In vivo*, EIF4E1 has lower abundance and cap-binding affinity than EIF4E3 and EIF4E4 (55,63), so EIF4E1 probably could not compete with EIF4E3 or EIF4E4 for cap binding by itself. Under the model in 9A, 4EIP would prevent the interaction of EIF4E1 with EIF3A, in partial analogy to Opisthokont 4E-BPs. An alternative is that EIF4E1-4EIP complex is functionally similar to 4EHP-GYF2 and that EIF4E1 acts as a constitutively inhibitory translation factor (Fig 9B), with mRNA-binding enhanced by cooperative interactions with 4EIP. If the 4EIP interaction prevents cap binding by EIF4E1 (65), this might allow decapping (Fig 9C).

**Figure 9.**
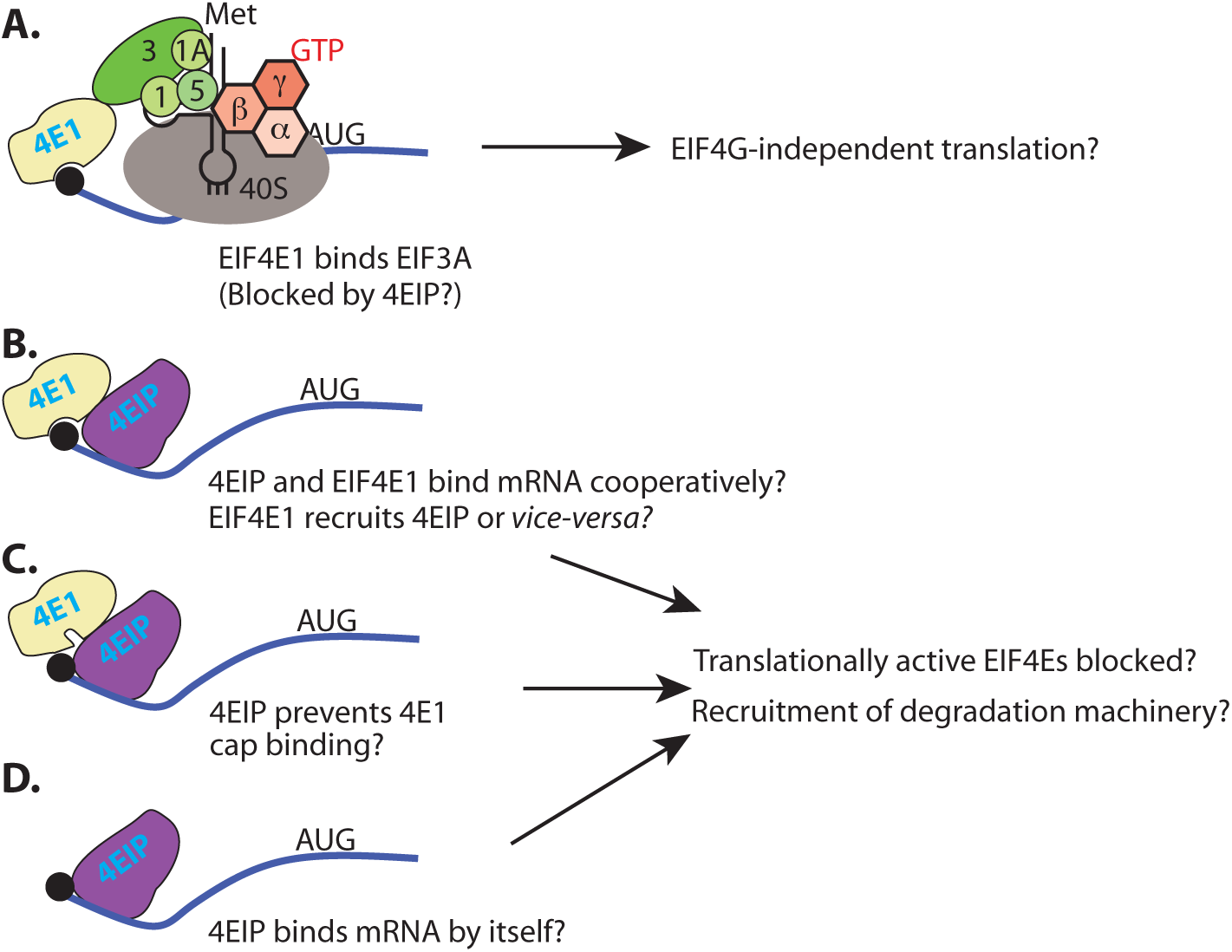
Models for EIF4E1 and 4EIP function. For details see the Discussion section.

The models in Figs 9B and C imply that the EIF4E1-4EIP complex prevents binding of active cap-binding initiation complexes. This is very likely to be the major effect in growing intact cells. However, several results indicated that the suppressive activity of 4EIP does not depend on its interaction with EIF4E1. First, both ∆N4EIP, which does not detectably interact with EIF4E1, and a C-terminal fragment which lacks all possible interaction surfaces, were sufficient to repress expression when tethered to a reporter mRNA. Here, one could argue that this is an artefact of tethering, and that normally, 4EIP is recruited to the mRNA via the cap-binding activity of EIF4E1. Stronger evidence comes from the ability of ∆N4EIP to complement the differentiation defect of cells lacking full-length 4EIP (Fig 7C). This suggests that during stumpy formation 4EIP might act independently of EIF4E1 (Fig 9D). Such activity would imply that 4EIP has intrinsic RNA-binding activity. There is indeed some evidence for this. Lueong *et al.* (68) subjected bloodstream-form trypanosomes to UV in order to create protein-RNA cross-links, then purified mRNA under highly denaturing conditions, in order to detect only those proteins that were cross-linked to mRNA. 4EIP was reproducibly detected in the resulting mass spectrometry analyses (68) (Supplementary Table S1, sheet 1).

Our interaction studies provided no insights into the mechanism by which 4EIP suppresses translation and/or causes mRNA destruction. Apart from the 4E interaction motif, there is a second highly conserved charged domain which might be implicated in protein-protein or protein-RNA interactions. However when tethered, the highly unstructured and proline-glutamine-rich C-terminus was sufficient for activity. 4EIPs from several other kinetoplastids have polyglutamine repeats (Supplementary Fig S1). Such repeats are present in many regulatory RNA-binding proteins (68), including Bicoid and GIGYF2. These and other low-complexity regions have been implicated in protein aggregation, and in formation of hydrogels and RNA-protein granules (81). EIF4E1 is found in starvation granules (82) (http://tryptag.org/?query=EIF4E1), but it is distributed throughout the cytosol and nucleus under normal conditions (55).

Both the absence of 4EIP in bloodstream forms, and its over-expression (as a fusion protein) in procyclic forms, resulted in a slight growth defect. We therefore suggest that in growing cells, 4EIP may help to suppress expression of proteins that are needed in only modest amounts. Upon over-expression, the suppression of genuine targets may be excessive, but in addition, 4EIP may bind to mRNAs with which it does not normally associate. (Whether this would require interaction with EIF4E1, or whether EIF4E1 levels would be limiting, is an open question since the abundance of 4EIP is not known.) When pleomorphic trypanosomes were incubated for prolonged periods at high density, the level of 4EIP became critical. Over-expression of 4EIP enhanced parasite survival, suggesting that translational shut-down is protective. In contrast, cells that lacked 4EIP showed a minor delay in suppression of protein synthesis. A common mechanism for general translation suppression is phosphorylation of EIF2I at threonine 169, but this is known not to be required for differentiation (83). The loss in translation in stumpy forms is therefore likely to be due primarily to loss of mRNA, secondary to transcription arrest. Despite the relatively mild effect on translation, however, trypanosomes lacking 4EIP were unable to convert to growing procyclic forms. Repression by 4EIP is therefore an obligatory part of the developmental programme.

## Methods

### Trypanosomes and plasmids

All experiments apart from those concerning differentiation were done using Lister 427 strain bloodstream forms or procyclic forms. For differentiation experiments, the EATRO1125 strain was used. The EATRO1125 bloodstream forms were routinely grown in the presence of 1.1% methyl cellulose (79) to retain differentiation capability. Otherwise, transfection and growth conditions were as described (73,84). Plasmids and oligonucleotides are listed in Supplementary Table S4. All growth and differentiation experiments were done in the absence of selecting drugs, and with at least a 24h prior pre-incubation without selection. Double knock-out lines were routinely grown without blasticidin and puromycin.

EATRO 1125 cells were grown at 37 °C, 5% carbon dioxide, in HMI-9 media containing 1.1 % methylcellulose. To obtain high density EATRO 1125 and stumpy cells, 5×10^5^ cells were allowed to grow to maximum density. Collection was as described in (79): cultures were diluted 5 times with PBS, filtered through 615 ¼ MN filters (Macherey-Nagel, Germany) and centrifuged at 1218 × g for 10 min. This collection procedure lasted up to 20 min.

### Polysomes, RNA, RNASeq and Northern blotting

Polysome analysis and Northern blotting were done as described in (72). For RNA preparation from differentiating cells, ~5×10^6^ cells (0 hr time point) ~3×10^7^ cells (36 hr time point) or ~5×10^7^ cells (48 hr time point) were collected. This yield at least 1.5μg of RNA, but the RNA from the 36h and 48h was degraded so could not be evaluated.

To find mRNAs that were bound to 4EIP, lysates from cells inducibly expressing 4EIP-TAP (not irradiated) were bound to IgG beads and washed. RNA was released from the beads by protease treatment, and purified. Input and unbound fractions were rRNA-depleted using oligonucleotides and RNaseH, all fractions were subjected to RNASeq (E-MTAB-6240), and the data were analysed, as described previously (73,85,86).

### Immunoprecipitation, Western blotting, tethering and interactome analysis

Immunoprecipitation, yeast two-hybrid screening, tethering assays and tandem affinity purification were all done as described in (53). Western blots were detected as described in (73). Apart from commercial antibodies to myc and V5, which came from various sources, antibodies were to ribosomal protein S9 (this lab), PAD1 (Keith Matthews, University of Edinburgh) and trypanothione reductase (Luise Krauth-Siegel, Heidelberg University).

### Pulse-labelling

For each time point ~3×10^7^ cells were collected, washed twice with PBS and re-suspended in 350μl labelling medium (Dulbecco’s Modified Eagle Medium (gibco) lacking L-methionine). These were grown in the conditions mentioned above for 1 hour before addition of ^35^S- methionine for 20 min. Thereafter the cells were collected by centrifugation at 3500 ×g for 2 min, washed twice with PBS and re-suspended in laemmli buffer.

### Availability

The ArrayExpress accession number for the 4EIP-TAP pull-down is E-MTAB-6240.

## Acknowledgements

We thank Thomas Ruppert and the Mass Spectrometry facility at the ZMBH for mass spectrometry analysis, and David Ibbersson (Bioquant, Heidelberg) for building RNASeq libraries and taking them for sequencing. We acknowledge contributions from MSc or BSc students during lab practicals, especially Jacob Borold who did preliminary polysome analysis after tethering. We thank Keith Matthews for antibody to PAD1 and Luise Krauth-Siegel for antibody to trypanothione reductase. We also thank Michal Shapira, Osvaldo de Melo Neto and Eden Freire for useful discussions and for communicating unpublished results. This work was supported by various grants from the Deutsche Forschungsgemeinschaft, including Cl112-17-1, Cl112-17-2, Cl112-24, and Cl112-28. Kevin Kamanyi is supported by a fellowship from the DAAD, and Franziska Egler has a stipend from HBIGS (Heidelberg Biosciences International Graduate School).

Author contributions were as follows:

M. Terrao: 4EIP-TAP experiments (Fig S3, Table S1), RNA pull-down including data analysis (Fig 5, Table S3), and the 2-hybrid screen (Table S2).

K. Kamanyi: Differentiation experiments, add-backs, tethering of EIF4E1 in 4EIP knockout. Figs 1E, 3, 6, 7, S7, S8, S9.

D. Droll: Figs 1A-D, S2, S3.

I. Minia: Fig 4.

E. Mugo: tethering in bloodstream forms (Fig 3) and RNAi in procyclics.

F. Egler: EIF4E1 DKO, Fig 8 and S10

J. Braun: 4EIP KO in Lister427 and EATRO1125, figs 6, S7

C. Clayton: Data analysis, funding acquisition, first draft of paper.

